# Mass spectrometry-based *de novo* sequencing of the anti-FLAG-M2 antibody using multiple proteases and a dual fragmentation scheme

**DOI:** 10.1101/2021.01.07.425675

**Authors:** Weiwei Peng, Matti F. Pronker, Joost Snijder

**Author notes:** equal contribution (Weiwei Peng, Matti F. Pronker).

## Abstract

Antibody sequence information is crucial to understanding the structural basis for antigen binding and enables the use of antibodies as therapeutics and research tools. Here we demonstrate a method for direct *de novo* sequencing of monoclonal IgG from the purified antibody products. The method uses a panel of multiple complementary proteases to generate suitable peptides for *de novo* sequencing by LC-MS/MS in a bottom-up fashion. Furthermore, we apply a dual fragmentation scheme, using both stepped high-energy collision dissociation (stepped HCD) and electron transfer high-energy collision dissociation (EThcD) on all peptide precursors. The method achieves full sequence coverage of the monoclonal antibody Herceptin, with an accuracy of 99% in the variable regions. We applied the method to sequence the widely used anti-FLAG™-M2 mouse monoclonal antibody, which we successfully validated by remodeling a high-resolution crystal structure of the Fab and demonstrating binding to a FLAG™-tagged target protein in Western blot analysis. The method thus offers robust and reliable sequences of monoclonal antibodies.

## Introduction

Antibodies can bind a great molecular diversity of antigens, owing to the high degree of sequence diversity that is available through somatic recombination, hypermutation, and heavy-light chain pairings ^1–2^. Sequence information on antibodies therefore is crucial to understanding the structural basis of antigen binding, how somatic hypermutation governs affinity maturation, and an overall understanding of the adaptive immune response in health and disease, by mapping out the antibody repertoire. Moreover, antibodies have become invaluable research tools in the life sciences and ever more widely developed as therapeutic agents ^3–4^. In this context, sequence information is crucial for the use, production and validation of these important research tools and biopharmaceutical agents ^5–6^.

Antibody sequences are typically obtained through cloning and sequencing of the coding mRNAs of the paired heavy and light chains ^7–9^. The sequencing workflows thereby rely on isolation of the antibody-producing cells from peripheral blood monocytes, or spleen and bone marrow tissues. These antibody-producing cells are not always readily available however, and cloning/sequencing of the paired heavy and light chains is a non-trivial task with a limited success rate ^7–9^. Moreover, antibodies are secreted in bodily fluids and mucus. Antibodies are thereby in large part functionally disconnected from their producing B-cell, which raises questions on how the secreted antibody pool relates quantitatively to the underlying B-cell population and whether there are potential sampling biases in current antibody sequencing strategies.

Direct mass spectrometry (MS)-based sequencing of the secreted antibody products is a useful complementary tool that can address some of the challenges faced by conventional sequencing strategies relying on cloning/sequencing of the coding mRNAs ^10–17^. MS-based methods do not rely on the availability of the antibody-producing cells, but rather target the polypeptide products directly, offering the prospect of a next generation of serology, in which secreted antibody sequences might be obtained from any bodily fluid. Whereas MS-based *de novo* sequencing still has a long way to go towards this goal, owing to limitations in sample requirements, sequencing accuracy, read length and sequence assembly, MS has been successfully used to profile the antibody repertoire and obtain (partial) antibody sequences beyond those available from conventional sequencing strategies based on cloning/sequencing of the coding mRNAs ^10–17^.

Most MS-based strategies for antibody sequencing rely on a proteomics-type bottom-up LC-MS/MS workflow, in which the antibody product is digested into smaller peptides for MS analysis ^14, 18–23^. Available germline antibody sequences are then often used either as a template to guide assembly of *de novo* peptide reads (such as in PEAKS Ab) ^23^, or used as a starting point to iteratively identify somatic mutations to arrive at the mature antibody sequence (such as in Supernovo) ^21^. To maximize sequence coverage and aid read assembly, these MS-based workflows typically use a combination of complementary proteases and aspecific digestion to generate overlapping peptides. The most straightforward application of these MS-based sequencing workflows is the successful sequencing of monoclonal antibodies from (lost) hybridoma cell lines, but it also forms the basis of more advanced and challenging applications to characterize polyclonal antibody mixtures and profile the full antibody repertoire from serum.

Here we describe an efficient protocol for MS-based sequencing of monoclonal antibodies. The protocol requires approximately 200 picomol of the antibody product and sample preparation can be completed within one working day. We selected a panel of 9 proteases with complementary specificities, which are active in the same buffer conditions for parallel digestion of the antibodies. We developed a dual fragmentation strategy for MS/MS analysis of the resulting peptides to yield rich sequence information from the fragmentation spectra of the peptides. The protocol yields full and deep sequence coverage of the variable domains of both heavy and light chains as demonstrated on the monoclonal antibody Herceptin. As a test case, we used our protocol to sequence the widely used anti-FLAG™-M2 mouse monoclonal antibody, for which no sequence was publicly available despite its described use in 5000+ peer-reviewed publications ^24–25^. The protocol achieved full sequence coverage of the variable domains of both heavy and light chains, including all complementarity determining regions (CDRs). The obtained sequence was successfully validated by remodeling the published crystal structure of the anti-FLAG™-M2 Fab and demonstrating binding of the synthetic recombinant antibody following the experimental sequence to a FLAG™-tagged protein in Western blot analysis. The protocol developed here thus offers robust and reliable sequencing of monoclonal antibodies with prospective applications for sequencing secreted antibodies from bodily fluids.

## Results

We used an in-solution digestion protocol, with sodium-deoxycholate as the denaturing agent, to generate peptides from the antibodies for LC-MS/MS analysis. Following heat denaturation and disulfide bond reduction, we used iodoacetic acid as the alkylating agent to cap free cysteines. Note that conventional alkylating agents like iodo-/chloroacetamide generate +57 Da mass differences on cysteines and primary amines, which may lead to spurious assignments as glycine residues in *de novo* sequencing. The +58 Da mass differences generated by alkylation with iodoacetic acid circumvents this potential pitfall.

We chose a panel of 9 proteases with activity at pH 7.5-8.5, so that the denatured, reduced and alkylated antibodies could be easily split for parallel digestion under the same buffer conditions. These proteases (with indicated cleavage specificities) included: trypsin (C-terminal of R/K), chymotrypsin (C-terminal of F/Y/W/M/L), α-lytic protease (C-terminal of T/A/S/V), elastase (unspecific), thermolysin (unspecific), lysN (N-terminal of K), lysC (C-terminal of K), aspN (N-terminal of D/E), and gluC (C-terminal of D/E). Correct placement or assembly of peptide reads is a common challenge in *de novo* sequencing, which can be facilitated by sufficient overlap between the peptide reads. This favors the occurrence of missed cleavages and longer reads, so we opted to perform a brief 4-hour digestion. Following digestion, SDC is removed by precipitation and the peptide supernatant is desalted, ready for LC-MS/MS analysis. The resulting raw data was used for automated *de novo* sequencing with the Supernovo software package.

As peptide fragmentation is dependent on many factors like length, charge state, composition and sequence ^26^, we needed a versatile fragmentation strategy to accommodate the diversity of antibody-derived peptides generated by the 9 proteases. We opted for a dual fragmentation scheme that applies both stepped high-energy collision dissociation (stepped HCD) and electron transfer high-energy collision dissociation (EThcD) on all peptide precursors ^27–29^. The stepped HCD fragmentation includes three collision energies to cover multiple dissociation regimes and the EThcD fragmentation works especially well for higher charge states, also adding complementary c/z ions for maximum sequence coverage.

We used the monoclonal antibody Herceptin (also known as Trastuzumab) as a benchmark to test our protocol ^30–31^. From the total dataset of 9 proteases, we collected 4408 peptide reads (defined as peptides with score >=500, see methods for details), 2866 of which with superior stepped HCD fragmentation, and 1722 with superior EThcD fragmentation (see Table S1). Sequence coverage was 100% in both heavy and light chains across the variable and constant domains (see Figures S1 and S2). The median depth of coverage was 148 overall and slightly higher in the light chain (see Table S1 and Figure S1-2). The median depth of coverage in the CDRs of both chains ranged from 42 to 210.

The experimentally determined *de novo* sequence is shown alongside the known Herceptin sequence for the variable domains of both chains in Figure 1, with exemplary MS/MS spectra for the CDRs. We achieved an overall sequence accuracy of 99% with the automated sequencing procedure of Supernovo, with 3 incorrect assignments in the light chain. In framework 3 of the light chain, I75 was incorrectly assigned as the isomer Leucine (L), a common MS-based sequencing error. In CDRL3 of the light chain, an additional misassignment was made for the dipeptide H91/Y92, which was incorrectly assigned as W91/N92. The dipeptides HY and WN have identical masses, and the misassignment of W91/N92 (especially W91) was poorly supported by the fragmentation spectra, in contrast to the correct H91/Y92 assignment (see c6/c7 in fragmentation spectra, Figure 1). Overall, the protocol yielded highly accurate sequences at a combined 230/233 positions of the variable domains in Herceptin.

**Figure 1.**
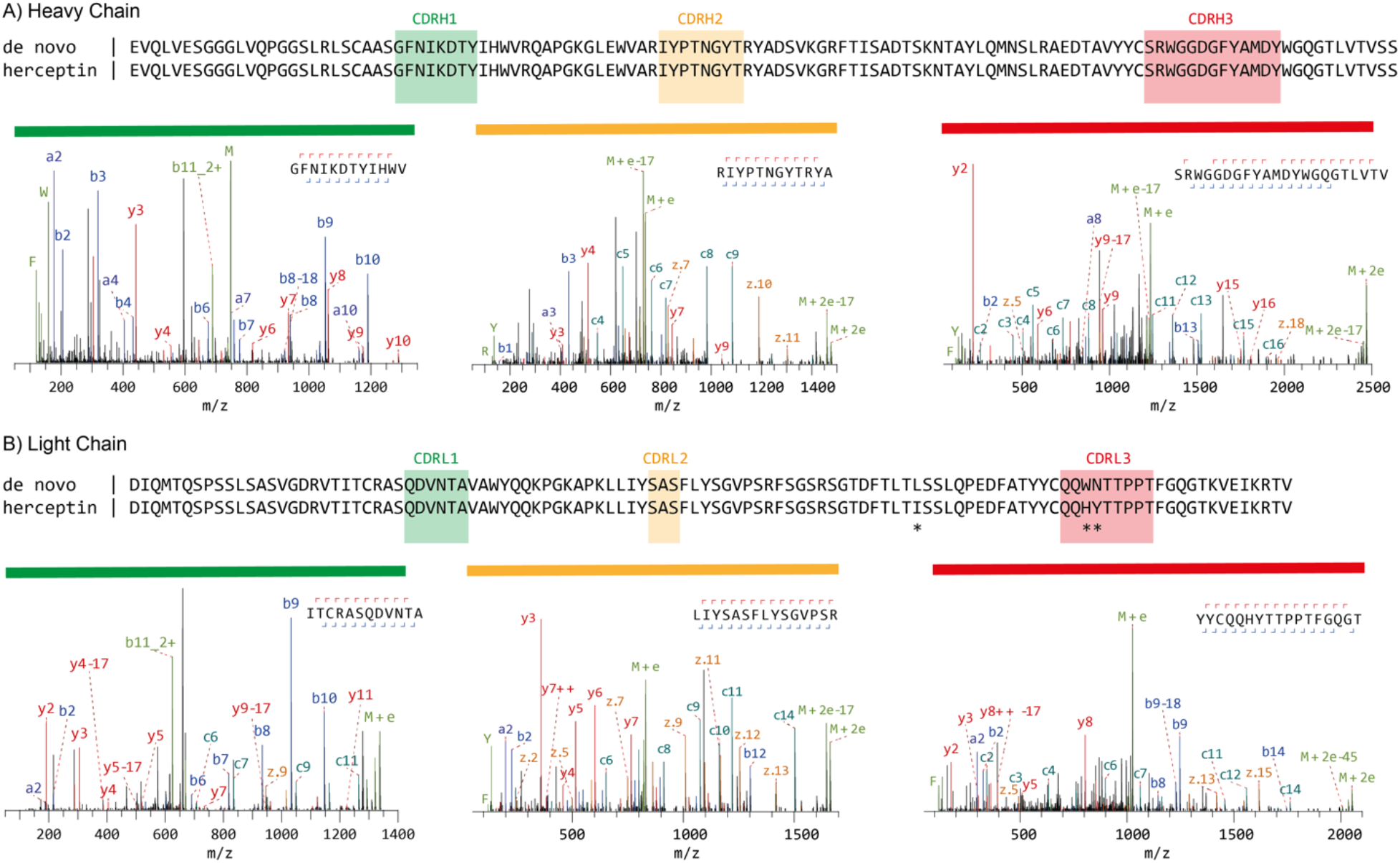
mass spectrometry-based *de novo* sequencing of the monoclonal antibody Herceptin. The variable regions of the Heavy (A) and Light Chains (B) are shown. The MS-based sequence is shown alongside the known Herceptin sequence, with differences highlighted by asterisks (*). Exemplary MS/MS spectra supporting the assigned sequences of the Heavy and Light Chain CDRs are shown below the alignments. Peptide sequence and fragment coverage are indicated on top of the spectra, with b/c ions indicated in blue and y/z ions in red. The same coloring is used to annotate peaks in the spectra, with additional peaks such as intact/charge reduced precursors, neutral losses and immonium ions indicated in green. Note that to prevent overlapping peak labels, only a subset of successfully matched peaks is annotated.

We next applied our sequencing protocol to the mouse monoclonal anti-FLAG™-M2 antibody as a test case ^24^. Despite the widespread use of anti-FLAG™-M2 to detect and purify FLAG™-tagged proteins ^32^, the only publicly available sequences can be found in the crystal structure of the Fab ^33^. The modelled sequence of the original crystal structure had to be inferred from germline sequences that could match the experimental electron density and also includes many placeholder Alanines at positions that could not be straightforwardly interpreted. The full anti-FLAG™-M2 dataset from the 9 proteases included 3371 peptide reads (with scores >= 500); 1983 with superior stepped HCD fragmentation spectra, and 1388 with superior EThcD spectra. We achieved full sequence coverage of the variable regions of both heavy and light chains, with a median depth of coverage in the CDRs ranging from 32 to 192 (see Table S1). As for Herceptin, the depth of coverage was better in the light chain compared to the heavy chain (see Figure S1-S2). The full MS-based anti-FLAG™-M2 sequences can be found in FASTA format in the supplementary information.

The MS-based sequences of anti-FLAG™-M2 are shown alongside the crystal structure sequences and the inferred germline precursors with exemplary MS/MS spectra for the CDRs in Figure 2. The experimentally determined sequence reveals that anti-FLAG™-M2 is a mouse IgG1, with an IGHV1-04/IGHJ2 heavy chain and IGKV1-117/IGKJ1 kappa light chain. The experimentally determined sequence differs at 34 and 9 positions in the heavy and light chain of the Fab crystal structure, respectively. To validate the experimentally determined sequences, we remodeled the crystal structure using the MS-based heavy and light chains, resulting in much improved model statistics (see Figure 3 and Table S2). The experimental electron densities show excellent support of the MS-based sequence (as shown for the CDRs in Figure 3B). A notable exception is L51 in CDRH2 of the heavy chain. The MS-based sequence was assigned as Leucine, but the experimental electron density supports assignment of the isomer Isoleucine instead (see Figure S3). In contrast to the original model our new MS-based model reveals a predominantly positively charged paratope (see Figure S4), which potentially complements the −3 net charge of the FLAG™ tag epitope (DYKDDDDK) to mediate binding. The experimentally determined anti-FLAG™-M2 sequence, with the L51I correction, was further validated by testing binding of the synthetic recombinant antibody to a purified FLAG™-tagged protein in Western blot analysis (see Figure 3C and S5). The synthetic recombinant antibody showed equivalent binding compared to the original antibody sample used for sequencing, confirming that the experimentally determined sequence is reliable to obtain the recombinant antibody product with the desired functional profile.

**Figure 2.**
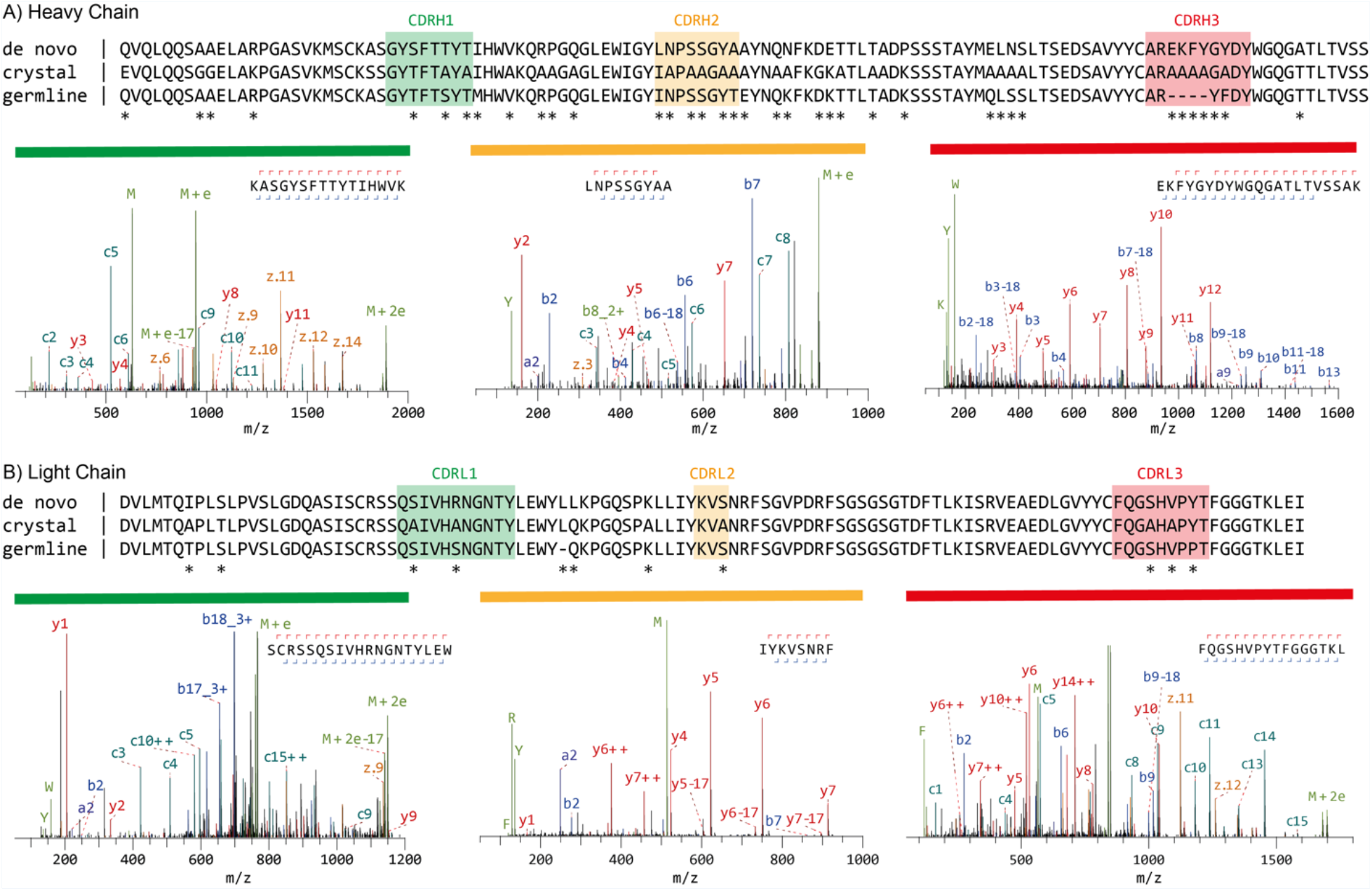
Mass spectrometry based *de novo* sequence of the mouse monoclonal anti-FLAG™-M2 antibody. The variable regions of the Heavy (A) and Light Chains (B) are shown. The MS-based sequence is shown alongside the previously published sequenced in the crystal structure of the Fab (PDB ID: 2G60), and germline sequence (IMGT-DomainGapAlign; IGHV1-04/IGHJ2; IGKV1-117/IGKJ1). Differential residues are highlighted by asterisks (*). Exemplary MS/MS spectra in support of the assigned sequences are shown below the alignments. Peptide sequence and fragment coverage are indicated on top of the spectra, with b/c ions indicated in blue, y/z ions in red. The same coloring is used to annotate peaks in the spectra, with additional peaks such as intact/charge reduced precursors, neutral losses and immonium ions indicated in green. Note that to prevent overlapping peak labels, only a subset of successfully matched peaks is annotated.

**Figure 3.**
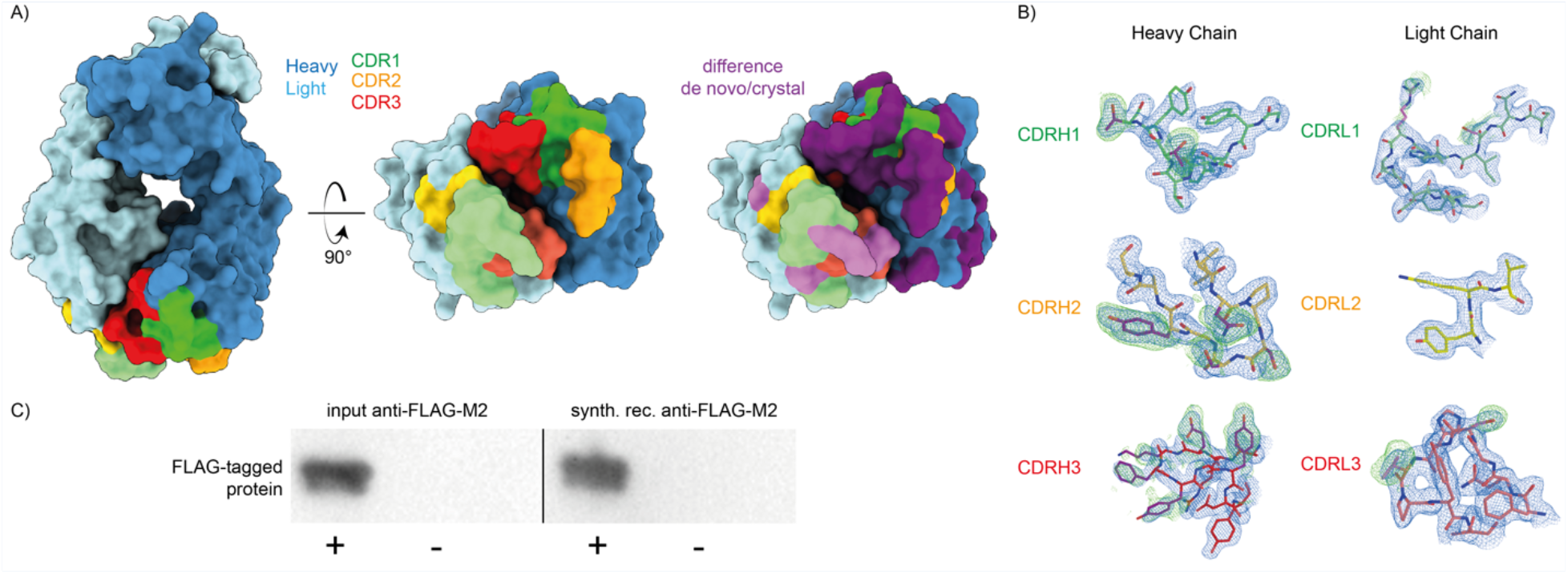
Validation of the MS-based anti-FLAG™-M2 sequence. A) the previously published crystal structure of the anti-FLAG™-M2 FAb was remodeled with the experimentally determined sequence, shown in surface rendering with CDRs and differential residues highlighted in colors. B) 2Fo-Fc electron density of the new refined map contoured at 1 RMSD is shown in blue and Fo-Fc positive difference density of the original deposited map contoured at 1.7 RMSD in green around the CDR loops of the heavy and light chains. Differential residues between the published crystal structure and the model based on our antibody sequencing are indicated in purple. C) Western blot validation of the synthetic recombinant anti-FLAG™-M2 antibody produced with the experimentally determined sequence demonstrate equivalent FLAG™-tag binding compared to commercial anti-FLAG™-M2 (see also Figure S3).

## Discussion

There are four other monoclonal antibody sequences against the FLAG™ tag publicly available through the ABCD (AntiBodies Chemically Defined) database ^34–36^. Comparison of the CDRs of anti-FLAG™-M2 with these additional four monoclonal antibodies reveals a few common motifs that may determine FLAG™-tag binding specificity (see Table S3). In the heavy chain, the only common motif between all five monoclonals is that the first three residues of CDRH1 follow a GXS sequence. In addition, the last three residues of CDRH3 of anti-FLAG™-M2 are YDY, similar to MDY in 2H8, and YDF in EEh13.6 (and EEh14.3 also ends CDRH3 with an aromatic F residue). In contrast to the heavy chain, the CDRs of the light chain are almost completely conserved in 4/5 monoclonals with only minimal differences compared to germline. The anti-FLAG™-M2 and 2H8 monoclonals were specifically raised in mice against the FLAG™-tag epitope ^24, 35^, whereas the computationally designed EEh13.6 and EEh14.3 monoclonals contain the same light chain from an EE-dipeptide tag directed antibody ^34^. This suggests that the IGKV1-117/IGKJ1 light chain may be a common determinant of binding to a small negatively charged peptide epitope like the FLAG™-tag and is readily available as a hardcoded germline sequence in the mouse antibody repertoire.

The availability of the anti-FLAG™-M2 sequences may contribute to the wider use of this important research tool, as well as the development and engineering of better FLAG™-tag directed antibodies. This example illustrates that our MS-based sequencing protocol yields robust and reliable monoclonal antibody sequences. The protocol described here also formed the basis of a recent application where we sequenced an antibody directly from patient-derived serum, using a combination with top-down fragmentation of the isolated Fab fragment ^37^. The dual fragmentation strategy yields high-quality spectra suitable for *de novo* sequencing and may further contribute to the exciting prospect of a new era of serology in which antibody sequences can be directly obtained from bodily fluids.

## Experimental Section

### Sample preparation

Anti-FLAG™ M2 antibody was purchased from Sigma (catalogue number F1804). Herceptin was provided by Roche (Penzberg, Germany). 27 μg of each sample was denatured in 2% sodium deoxycholate (SDC), 200 mM Tris-HCl, 10 mM tris(2-carboxyethyl)phosphine (TCEP), pH 8.0 at 95°C for 10 min, followed with 30 min incubation at 37°C for reduction. Sample was then alkylated by adding iodoacetic acid to a final concentration of 40 mM and incubated in the dark at room temperature for 45 min. 3 μg Sample was then digested by one of the following proteases: trypsin, chymotrypsin, lysN, lysC, gluC, aspN, aLP, thermolysin and elastase in a 1:50 ratio (w:w) in a total volume of 100 uL of 50 mM ammonium bicarbonate at 37°C for 4 h. After digestion, SDC was removed by adding 2 uL formic acid (FA) and centrifugation at 14000 g for 20 min. Following centrifugation, the supernatant containing the peptides was collected for desalting on a 30 µm Oasis HLB 96-well plate (Waters). The Oasis HLB sorbent was activated with 100% acetonitrile and subsequently equilibrated with 10% formic acid in water. Next, peptides were bound to the sorbent, washed twice with 10% formic acid in water and eluted with 100 µL of 50% acetonitrile/5% formic acid in water (v/v). The eluted peptides were vacuum-dried and reconstituted in 100 µL 2% FA.

### Mass Spectrometry

The digested peptides (single injection of 0.2 ug) were separated by online reversed phase chromatography on an Agilent 1290 UHPLC (column packed with Poroshell 120 EC C18; dimensions 50 cm x 75 µm, 2.7 µm, Agilent Technologies) coupled to a Thermo Scientific Orbitrap Fusion mass spectrometer. Samples were eluted over a 90 min gradient from 0% to 35% acetonitrile at a flow rate of 0.3 μL/min. Peptides were analyzed with a resolution setting of 60000 in MS1. MS1 scans were obtained with standard AGC target, maximum injection time of 50 ms, and scan range 350-2000. The precursors were selected with a 3 m/z window and fragmented by stepped HCD as well as EThcD. The stepped HCD fragmentation included steps of 25%, 35% and 50% NCE. EThcD fragmentation was performed with calibrated charge-dependent ETD parameters and 27% NCE supplemental activation. For both fragmentation types, ms2 scan were acquired at 30000 resolution, 800% Normalized AGC target, 250 ms maximum injection time, scan range 120-3500.

### MS Data Analysis

Automated *de novo* sequencing was performed with Supernovo (version 3.10, Protein Metrics Inc.). Custom parameters were used as follows: non-specific digestion; precursor and product mass tolerance was set to 12 ppm and 0.02 Da respectively; carboxymethylation (+58.005479) on cysteine was set as fixed modification; oxidation on methionine and tryptophan was set as variable common 1 modification; carboxymethylation on the N-terminus, pyroglutamic acid conversion of glutamine and glutamic acid on the N-terminus, deamidation on asparagine/glutamine were set as variable rare 1 modifications. Peptides were filtered for score >=500 for the final evaluation of spectrum quality and (depth of) coverage. Supernovo generates peptide groups for redundant MS/MS spectra, including also when stepped HCD and EThcD fragmentation on the same precursor both generate good peptide-spectrum matches. In these cases only the best-matched spectrum is counted as representative for that group. This criterium was used in counting the number of peptide reads reported in Table S1. Germline sequences and CDR boundaries were inferred using IMGT/DomainGapAlign ^38–39^.

### Revision of the anti-FLAG™-M2 Fab crystal structure model

As a starting point for model building, the reflection file and coordinates of the published anti-FLAG™-M2 Fab crystal structure were used (PDB ID: 2G60) ^33^. Care was taken to use the original *Rfree* labels of the deposited reflection file for refinement, so as not to introduce extra model bias. Differential residues between this structure and our mass spectrometry-derived anti-FLAG™ sequence were manually mutated and fitted in the density using Coot ^40^. Many spurious water molecules that caused severe steric clashes in the original model were also manually removed in Coot. Densities for two sulfate and one chloride ion were identified and built into the model. The original crystallization solution contained 0.1 M ammonium sulfate. Iterative cycles of model geometry optimization in real space in Coot and reciprocal space refinement by Phenix were used to generate the final model, which was validated with Molprobity ^41–42^.

### Cloning and expression of synthetic recombinant anti-FLAG™-M2

To recombinantly express full-length anti-FLAG™-M2, the proteomic sequences of both the light and heavy chains were reverse-translated and codon optimized for expression in human cells using the Integrated DNA Technologies (IDT) web tool (*http://www.idtdna.com/CodonOpt*) ^43^. For the linker and Fc region of the heavy chain, the standard mouse Ig gamma-1 (IGHG1) amino acid sequence (Uniprot P01868.1) was used. An N-terminal secretion signal peptide derived from human IgG light chain (MEAPAQLLFLLLLWLPDTTG) was added to the N-termini of both heavy and light chains. BamHI and NotI restriction sites were added to the 5’ and 3’ ends of the coding regions, respectively. Only for the light chain, a double stop codon was introduced at the 3’ site before the NotI restriction site. The coding regions were subcloned using BamHI and NotI restriction-ligation into a pRK5 expression vector with a C-terminal octahistidine tag between the NotI site and a double stop codon 3’ of the insert, so that only the heavy chain has a C-terminal AAAHHHHHHHH sequence for Nickel-affinity purification (the triple alanine resulting from the NotI site). The L51I correction in the heavy chain was introduced later (after observing it in the crystal structure) by IVA cloning ^44^. Expression plasmids for the heavy and light chain were mixed in a 1:1 (w/w) ratio for transient transfection in HEK293 cells with polyethylenimine, following standard procedures. Medium was collected 6 days after transfection and cells were spun down by 10 minutes of centrifugation at 1000*g*. Antibody was directly purified from the supernatant using Ni-sepharose excel resin (Cytiva Lifes Sciences), washing with 500 mM NaCl, 2 mM CaCl2, 15 mM imidazole, 20 mM HEPES pH 7.8 and eluting with 500 mM NaCl, 2 mM CaCl2, 200 mM imidazole, 20 mM HEPES pH 7.8.

### Western blot validation of anti-FLAG™-M2 binding

To test binding of our recombinant anti-FLAG™-M2 to the FLAG™-tag epitope, compared to the commercially available anti-FLAG™-M2 (Sigma), we used both antibodies to probe Western blots of a FLAG™-tagged protein in parallel. Purified Rabies virus glycoprotein ectodomain (SAD B19 strain, UNIPROT residues 20-450) with or without a C-terminal FLAG™-tag followed by a foldon trimerization domain and an octahistidine tag was heated to 95 °C in XT sample buffer (Biorad) for 5 minutes. Samples were run twice on a Criterion XT 4-12% polyacrylamide gel (Biorad) in MES XT buffer (Biorad) before Western blot transfer to a nitrocellulose membrane in tris-glycine buffer (Biorad) with 20% methanol. The membrane was blocked with 5% (w/v) dry non-fat milk in phosphate-buffered saline (PBS) overnight at 4 °C. The membrane was cut in two (one half for the commercial and one half for the recombinant anti-FLAG™-M2) and each half was probed with either commercial (Sigma) or recombinant anti-FLAG™-M2 at 1 µg/mL in PBS for 45 minutes. After washing three times with PBST (PBS with 0.1% v/v Tween20), polyclonal goat anti-mouse fused to horseradish peroxidase (HRP) was used to detect binding of anti-FLAG™-M2 to the FLAG™-tagged protein for both membranes. The membranes were washed three more times with PBST before applying enhanced chemiluminescence (ECL; Pierce) reagent to image the blots in parallel.

## Data Availability

The raw LC-MS/MS data have been deposited to the ProteomeXchange Consortium via the PRIDE partner repository with the dataset identifier PXD023419. The coordinates and reflection file with phases for the remodeled crystal structure of the anti-FLAG™-M2 Fab have been deposited in the Protein Data Bank under accession code 7BG1; as per wwPDB policy, the deposition will be released upon peer-reviewed publication of this work. The atomic coordinate file is therefore also available as supplementary information.

## Supporting information

Atomic Coordinates for Remodelled Fab (7BG1)

## Acknowledgements

Herceptin was a kind gift from Roche (Penzberg, Germany). We would like to acknowledge support by Protein Metrics Inc. through access to Supernovo software and helpful discussion on *de novo* antibody sequencing. We would like to thank everyone in the Biomolecular Mass Spectrometry and Proteomics group at Utrecht University for support and helpful discussions. This research was funded by the Dutch Research Council NWO Gravitation 2013 BOO, Institute for Chemical Immunology (ICI; 024.002.009).

## Author Contributions

WP and JS conceived of the project. WP carried out the MS experiments. WP and JS analyzed the MS data. MFP remodeled the crystal structure. MFP cloned and produced the synthetic recombinant antibody and carried out Western blotting. JS supervised the project. JS wrote the first draft and all authors contributed to preparing the final version of the manuscript.

## Supplementary Information

atomic coordinate file in .pdb format of remodeled Fab crystal structure

anti-FLAG™-M2 MS-based sequences in FASTA format

Tables S1-S3

Figures S1-S5

