## Supplementary Information for "Mass spectrometry-based *de novo* sequencing of the anti-FLAG-M2 antibody using multiple proteases and a dual fragmentation scheme"

[#]equal contribution

mass spectrometry, antibody, de novo sequencing, EThcD, stepped HCD, Herceptin, FLAG tag, anti-FLAG-M2.

**anti-FLAG-M2 MS-based sequence** (with L51I correction)

```
>anti-FLAG-M2_MS_HeavyChain
QVQLQQSAAELARPGASVKMSCKASGYSFTTYTIHWVKQRPGQGLEWIGYINPSSGYAAYNQNFKDETTLTADPSSS
TAYMELNSLTSEDSAVYYCAREKFYGYDYWGQGATLTVSSAKTTPPSVYPLAPGSAAQTNSMVTLGCLVKGYFPEPV
TVTWNSGSLSSGVHTFPAVLQSDLYTLSSSVTVPSSPRPSETVTCNVAHPASSTKVDKKIVPRDCGCKPCICTVPEV
SSVFIFPPKPKDVLTITLTPKVTCVVVDISKDDPEVQFSWFVDDVEVHTAQTQPREEQFNSTFRSVSELPIMHQDWL
NGKEFKCRVNSAAFPAPIEKTISKTKGRPKAPQVYTIPPPKEQMAKDKVSLTCMITDFFPEDITVEWQWNGQPAENY
KNTQPIMNTNGSYFVYSKLNVQKSNWEAGNTFTCSVLHEGLHNHHTEKSLSHSPGK

>anti-FLAG-M2_MS_LightChain
DVLMTQIPLSLPVSLGDQASISCRSSQSIVHRNGNTYLEWYLLKPGQSPKLLIYKVSNRFSGVPDRFSGSGSGTDFT
LKISRVEAEDLGVYYCFQGSHVPYTFGGGTKLEIRRADAAPTVSIFPPSSEQLTSGGASVVCFLNNFYPKDINVKWK
IDGSERQNGVLNSWTDQDSKDSTYSMSSTLTLTKDEYERHNSYTCEATHKTSTSPIVKSFNRNEC
```

**Table S1.** Coverage statistics for the Herceptin benchmark and anti-FLAG™-M2 MAb sequences.

| | | Herceptin | anti-FLAG-M2 |
|---|---|---|---|
| **# peptide reads** (Byonic score >=500) | total | 4408 | 3371 |
| | stepped HCD | 2686 | 1983 |
| | EThcD | 1722 | 1388 |
| **depth-of-coverage** (median [range]) | total | 148 [8-394] | 84 [0-382] |
| | **CDRH1** | 163 [158-176] | 32 [22-47] |
| | **CDRH2** | 94 [88-103] | 39 [36-43] |
| | **CDRH3** | 42 [18-67] | 66 [50-75] |
| | **CDRL1** | 210 [208-218] | 192 [144-207] |
| | **CDRL2** | 74 [71-84] | 46 [40-60] |
| | **CDRL3** | 140 [130-143] | 127 [109-131] |

**Table S2.** Model statistics for Fab crystal structure.

| Refinement statistics | | |
|---|---|---|
| Resolution (Å) | 42.52-1.86 | |
| No. of reflections | 39988 | |
| **PDB** | **2G60 (old)** | **7BG1 (new)** |
| Total number of atoms | 3518 | 3497 |
| Average atomic displacement parameter (Å$^2$) | 45.0 | 52.0 |
| $R_{work}/R_{free}$ | 0.235/0.278 | 0.217/0.255 |
| Bond length RMSZ | 0.93 | 0.28 |
| Bond angle RMSZ | 0.96 | 0.51 |
| Ramachandran favored/outliers (%) | 93.0/1.0 | 97.57/0.24 |
| Molprobity score | 3.37 | 1.60 |
| Clashscore | 56 | 3.61 |

**Table S3.** Comparison of CDR sequences from anti-FLAG™-M2 to other known FLAG™-tag binding MAbs (see refs 36-37).

| | Heavy Chain | | |
|---|---|---|---|
| **MAb** | **CDRH1** | **CDRH2** | **CDRH3** |
| anti-FLAG-M2 | GYSFTTYT---- | LNPSSGYA | AREKFYGYDY |
| 2H8 | GFSLNTSGRS-- | IYWDDDK | ARRMDY |
| EEh13.6 | GDSLSSFNAGVN | HGAVM-STR | AKSTGRYDF |
| EEh14.3 | GDSLSSYNAGVN | HMAGV-STR | VRNEWSGAF |
| EEf15.4 | GFSIK--GANVN | HVRGDASTR | ADRKMYSFYSGGEA |

| | Light Chain | | |
|---|---|---|---|
| **MAb** | **CDRL1** | **CDRL2** | **CDRL3** |
| anti-FLAG-M2 | QSIVHRNGNTY | KVS | FQGSHVPYT |
| 2H8 | QSLVHSNGNTY | KVS | SQSTHVPYT |
| EEh13.6 | QSIVHSNGNTY | KVS | FQGSLVPPT |
| EEh14.3 | QSIVHSNGNTY | KVS | FQGSLVPPT |
| EEf15.4 | NARSGS | DGN | SAFDQTNKYVG |

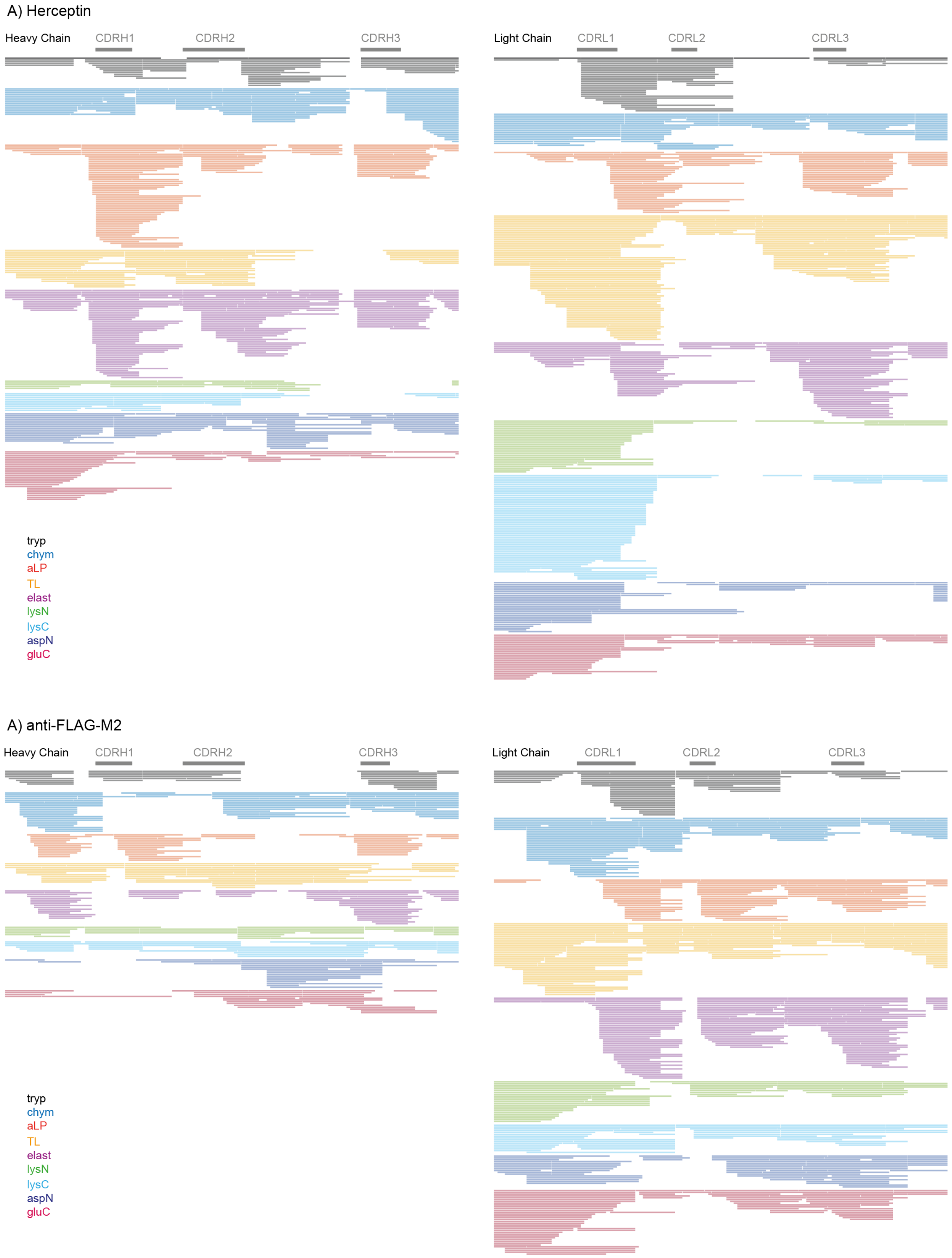


**Figure S1.** Coverage maps of Herceptin benchmark (A) and anti-FLAG™-M2 MAb (B) sequences. Peptides with Byonic scores of >=500 are shown.

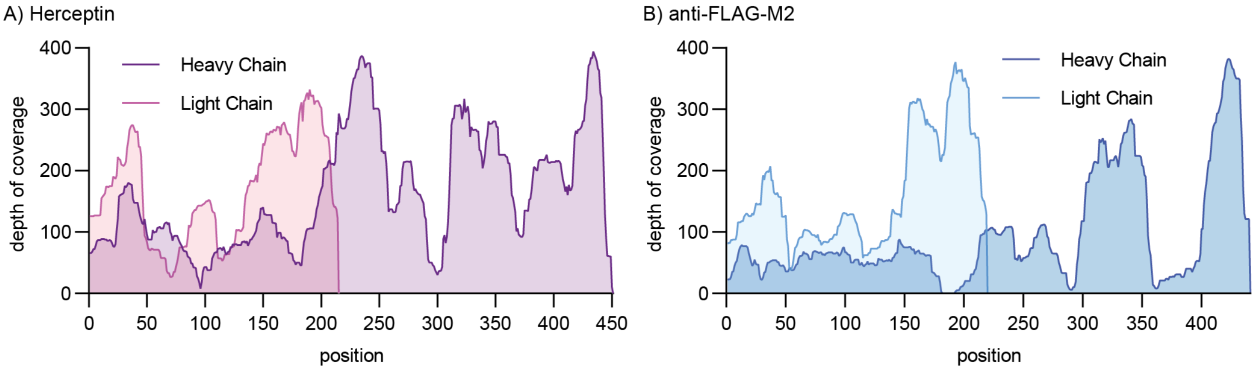


**Figure S2.** Depth of coverage profiles for Herceptin (A) and anti-FLAG™-M2 (B) sequences, based on peptides with Byonic score >=500, as in Figure S1.

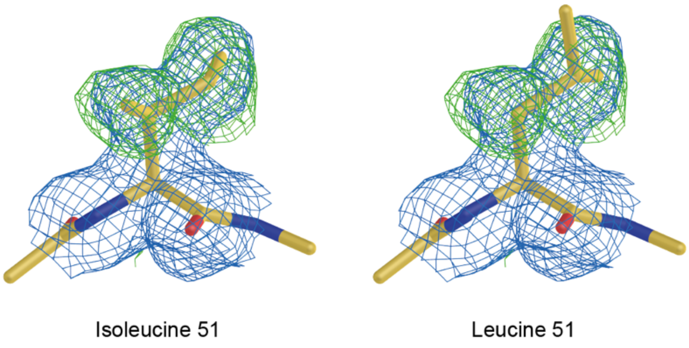


**Figure S3.** Isoleucine/Leucine assignment at Heavy Chain position 51 of anti-FLAG™-M2. (left panel) Electron density around isoleucine 51 at a contour level of 1.0 RMSD in blue and simulated annealing omit map density of the $C_{\gamma1}$, $C_{\gamma2}$ and $C_{\delta}$ atoms of this residue at a contour level of 2.5 R.M.S.D. in green. (right panel) A leucine instead of an isoleucine in this location has a poor fit to both maps.

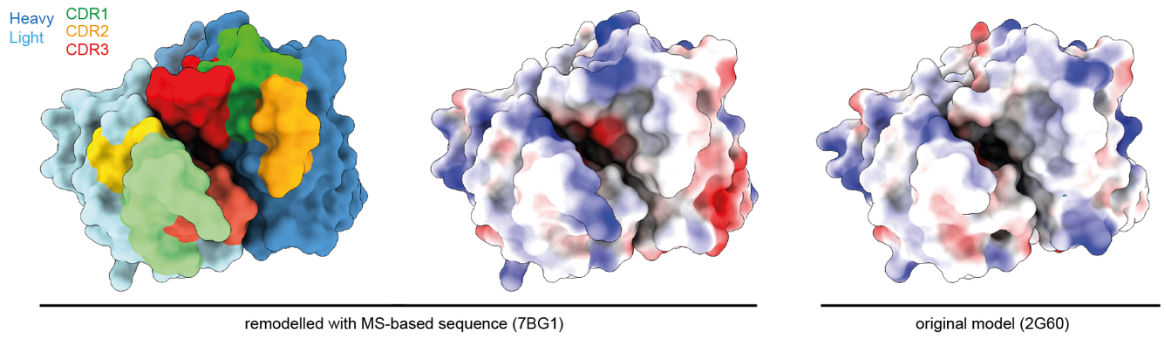


**Figure S4.** Electrostatic surface potential of the anti-FLAG™-M2 paratope. The revised crystal structure based on the MS-derived sequence (PDB ID: 7BG1) is shown alongside the original model (PDB ID: 2G60). The electrostatic surface was calculated with the default *coulombic* command in ChimeraX.

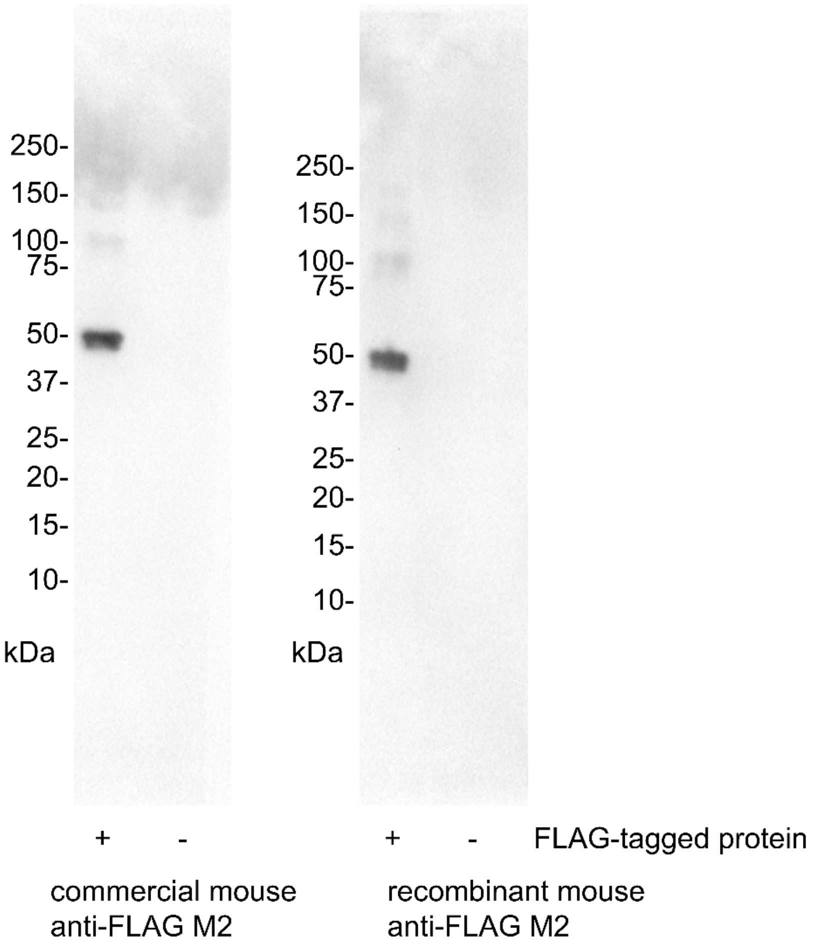


**Figure S5.** Western blot validation of synthetic recombinant anti-FLAG™-M2 compared to the originally sequenced sample. Same Western blot as shown in Figure 3C, showing complete lanes with marker positions.
